# Myocardial infarction creates a critical time window for AAV gene therapy

**DOI:** 10.1101/2024.06.10.597311

**Authors:** Gonglie Chen, Yueyang Zhang, Zhan Chen, Luzi Yang, Fei Gao, Erdan Dong, Yuxuan Guo

**Affiliations:** School of Basic Medical Sciences and Institute of Cardiovascular Sciences, State Key Laboratory of Vascular Homeostasis and Remodeling, Peking University Health Science Center, Beijing 100191, China; Department of Cardiology, Beijing Anzhen Hospital, Capital Medical University, Beijing 100029, China; Department of Cardiology and Institute of Vascular Medicine, Peking University Third Hospital, Beijing 100191, China; Research Center for Cardiopulmonary Rehabilitation, University of Health and Rehabilitation Sciences Qingdao Hospital (Qingdao Municipal Hospital), School of Health and Life Sciences, University of Health and Rehabilitation Sciences, Qingdao 266071, China; Beijing Key Laboratory of Cardiovascular Receptors Research, Beijing 100191, China

**Keywords:** Myocardial infarction, adeno-associated virus, gene delivery, gene therapy

## Abstract

Recombinant adeno-associated virus (AAV) is a major gene delivery vector for cardiac gene therapy. The factors that influence AAV-based cardiac gene transfer remain incompletely understood. This study showed that myocardial infarction (MI) enhanced cardiac AAV transduction and gene expression in mice after systemic administration, peaking at the third day post MI. These additional AAV vectors enriched at the infarcted region, correlated with the pathological permeabilization of the coronary vessels. The outcome of AAV-base gene therapy for MI, via *Camk2d* base editing, was significantly improved when AAV was injected at the third day post MI. Together, our findings uncovered a critical therapeutic time window after MI that facilitated AAV-based cardiac gene transfer, which could be harnessed to boost both basic and translational cardiology.

**Methods:** Adult C57BL/6 mice were subjected to left anterior descending (LAD) coronary artery ligation. Permanent LAD ligation created the MI model while 30min ligation followed by reperfusion established the ischemia/reperfusion (I/R) model. 2×10^11^ AAV vectors were injected into the mice via tail vein. The AAV vectors carried transgenes that were activated by the cardiomyocyte-specific *Tnnt2* promoter or the constitutively active CMV promoter. AAV expressed luciferase with the hemagglutinin (HA) tag. The amount of AAV vectors were quantified by real-time quantitative PCR (RT-qPCR) analysis of genomic DNA. Transgene expression was measured by RT-qPCR at the mRNA level or HA tag immunofluorescence imaging and western blot at the protein level. Luciferase activity was measured via bioluminescence imaging.

## Main text

Recombinant adeno-associated virus (AAV) is a popular gene delivery vector for gene therapy. A major obstacle in cardiac gene therapy involves the limited cardiac transduction efficiency by AAV. General strategies to enhance AAV gene transfer include capsid engineering, administration route improvement and neutralizing antibody management, but whether heart diseases exhibit unique properties that could be harnessed to boost AAV applications remain unclear.

Myocardial infarction (MI) is one of the leading causes of death and disability worldwide. Although an array of AAV gene therapy studies for MI have been conducted, a consensus on the AAV administration plan remains unestablished. Recently, several studies implicated an enhancing effect of myocardial ischemia on gene delivery vectors including not only AAV(1) but also lipid nanoparticles(2). Yet more extensive studies are necessary to determine its impacts on gene therapy outcomes.

To determine the effect of MI on AAV-based cardiac gene transfer, MI and sham mice were administered with 2×10^11^vg (vector genome) AAV serotype 9 (AAV9) carrying a HA-Luciferase transgene driven by the cardiac-specific Tnnt2 promoter at 3 days post infarction (Figure 1A). One week later, bioluminescence imaging demonstrated a robust increase of luciferase activity in MI hearts (Figure 1B). DNA and RNA were purified from the heart for real-time quantitative PCR (RT-qPCR). While vector DNA and its corresponding mRNA were both significantly higher in the MI group, the mRNA/DNA ratio, a metrics for promoter activity, exhibited no change (Figure 1C). Therefore, MI enhanced AAV gene expression by increasing the viral load in the heart.

**Figure 1.**
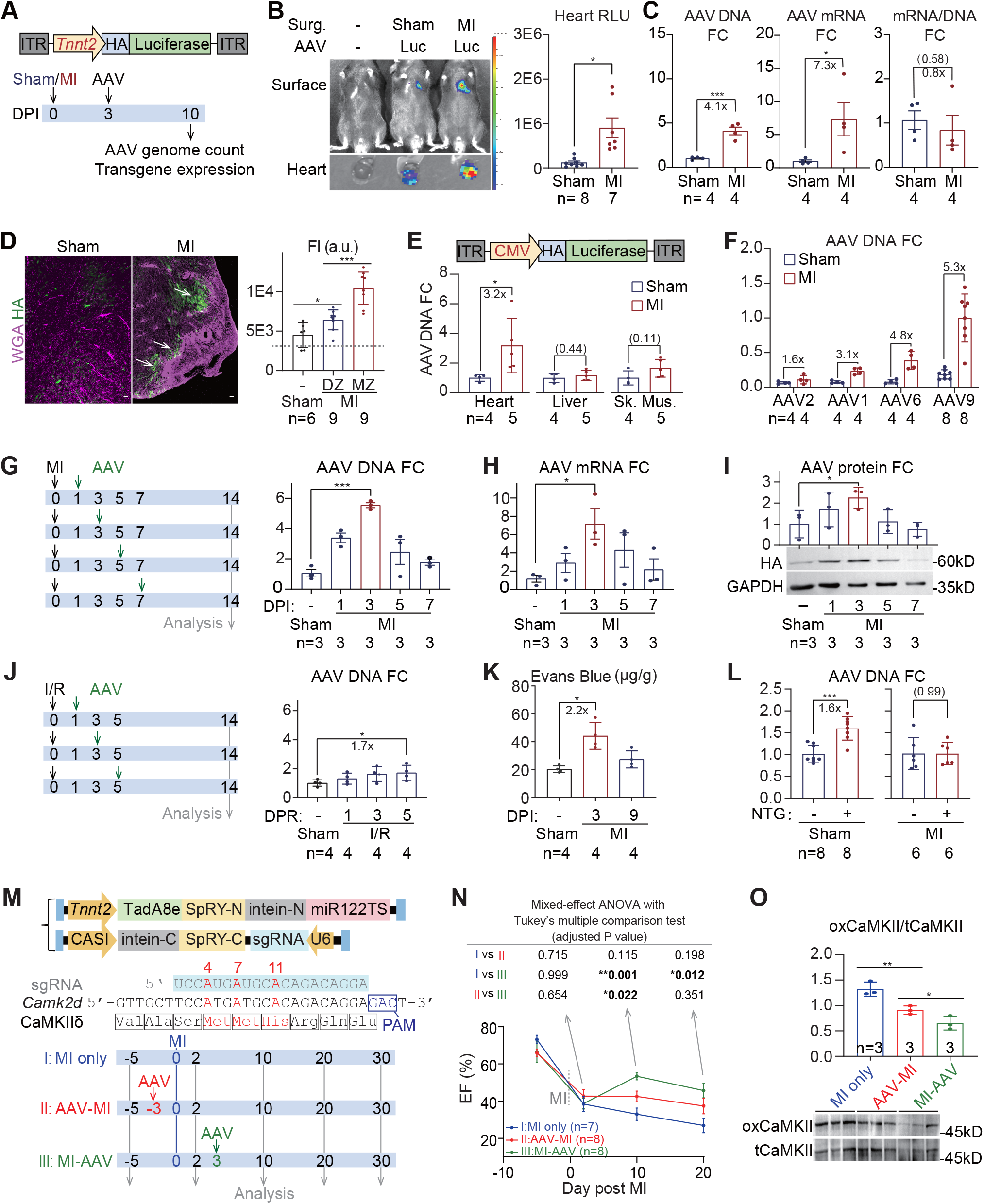
MI enhances AAV-based gene transfer to the heart. **(A)** A diagram of vector and experimental design. AAV, adeno-associated virus; ITR, inverted terminal repeats; HA, hemagglutinin tag; MI, myocardial infarction; DPI, day post MI. **(B)** Representative bioluminescence images (left) and quantification of signals (right) in the heart. Surg., surgery; Luc, luciferase; RLU, relative luminometer unit. **(C)** Real-time quantitative PCR (RT-qPCR) fold change (FC) analysis of AAV vector DNA (left), luciferase mRNA (middle) and mRNA/DNA (right) in the hearts. **(D)** Immunofluorescence images (left) of cardiac cryosections and quantification of fluorescence intensity (FI) of cardiac subdomains (right). Grey dashed line indicates mean background signal in non-AAV-treated control samples. Arrows point to zones with enhanced HA signals. Scale bar, 50 μm; a.u., arbitrary unit; MZ, marginal zone; DZ, distal zone. **(E)** A diagram of vector design and AAV DNA FC in variant organs. Sk. Mus., skeletal muscle. **(F)** vector DNA FC in hearts treated with various AAV serotypes. FCs are all relative to the AAV9-MI group. **(G-I)** A diagram of experimental design and analysis of vector DNA **(G)**, mRNA expression **(H)** and protein expression **(I)** in the heart when AAV was injected at serial time points. **(J)** A diagram of experimental design and RT-qPCR analysis of vector DNA in the heart. I/R, ischemia/reperfusion; DPR, day post reperfusion. **(K)** Quantification of Evans blue dye in myocardial extracts. 1% Evans blue was tail-vein injected at 4h before the dyes were extracted from the heart in formamide and quantified via 620nm absorption. **(L)** RT-qPCR analysis of AAV vector counts in sham vs. MI mice following 10 mg/kg NTG pre-treatment for 10 min. NTG, nitroglycerin. **(M)** AAV adenine base editing vectors with its target bases in red and an experimental design to compare preventative (AAV-MI) versus therapeutic (MI-AAV) AAV administration. **(N)** Echocardiogram and statistical analysis. EF, ejection fraction. **(O)** Western blot analysis of CaMKIIδ oxidation. oxCaMKII, oxidated CaMKII; tCaMKII, total CaMKII. All plots: mean ± SD; Welch’s t-test unless otherwise indicated: *P<0.05, **P<0.01, ***P<0.001, non-significant P value in parenthesis.

As an inroad to understand how MI hearts uptake more AAV than sham controls, immunostaining of HA-Luciferase was performed on cardiac cryosections. This experiment showed that MI elevated AAV expression preferentially at loci adjacent to the infarcted area (marginal zone, MZ), which were labelled by strong wheat germ agglutinin staining (Figure 1D, arrows). By contrast, distal zones (DZ) further away from the injured region exhibited less enhancement of the HA-Luciferase signal (Figure 1D). MI also enhanced the cardiac transduction of another AAV9-HA-Luciferase vector carrying the CMV promoter. No changes in AAV infection were detected in liver or skeletal muscle (Figure 1E). The enhancing effect of MI on cardiac gene transfer could also be observed when AAV1, AAV2 or AAV6 were applied (Figure 1F) although the AAV2<AAV1<AAV6<AAV9 rank of cardiac tropism in mice was retained(3). Therefore, MI enhanced AAV transduction independent of promoter or serotype.

AAV was next administered at serial time points after MI. While the viral DNA in the heart quickly increased when AAV was injected in the first three days after MI, it gradually declined afterwards and returned to the basal level by a week after MI (Figure 1G). MI maximally enhanced AAV transduction by 3-5 folds at day 3 post infarction (Figure 1C and G). This observation was also validated by RT-qPCR and western blot analysis of AAV expression in the heart (Figure 1H-I). In a model of cardiac ischemia-reperfusion injury (I/R), although I/R injury also moderately enhanced AAV transduction, this effect was much weaker than MI and no peak was detected at 3 days post reperfusion (Figure 1J). Thus, prolonged cardiac ischemia by MI was necessary to trigger the robust AAV-enhancing effect.

MI could locally disrupt the vascular barriers(4), enhancing the exposure of cardiomyocytes to AAV in the blood. To test this idea, an Evans Blue permeabilization assay was applied to confirm the temporarily enhanced vascular permeability at 3 days after MI (Figure 1K). Pretreatment of nitroglycerin (NTG), a classic vasodilatory drug that increases vascular permeability, increased cardiac AAV transduction in healthy hearts. However, this effect was diminished when NTG was applied at day 3 after MI (Figure 1L). These results suggested that MI enhanced cardiac AAV transduction partly by facilitating AAV passing through the vascular barrier, similar to the effect of NTG.

In gene therapy studies, early preventative AAV administration usually results in better outcomes than later treatments that attempt to reverse the disease phenotypes. However, the enhancing effect of MI on AAV transduction suggested the presence of a critical post-MI therapeutic window that could lead to better outcomes than preventative treatment. To test this hypothesis, we adopted a recently developed AAV base editing system that could ablate CaMKIIδ oxidation in the heart and slowed down the progression of MI-induced cardiac dysfunction(5). Specifically, 5×10^11^vg total AAV was injected via tail veins at 3 days either before or after MI induction to mimic the preventative versus therapeutic treatments, respectively (Figure M). While the therapeutic treatment temporally reversed the decline of ejection fraction (EF) in the first week after AAV injection, the beneficiary effect of the preventative administration on cardiac function appeared significantly weaker (Figure N). Western blot further validated the more efficient ablation of CaMKII oxidation in the therapeutic treatment group (Figure O).

Overall, this study indicates that MI creates a critical time window that facilitates AAV-mediated cardiac gene transfer. This effect could be harnessed to benefit both basic and translational cardiology relevant to the gene therapy for MI. Future mechanistic investigation of this phenomena is warranted to develop new AAV boosters similar to NTG as the key approach to enhance AAV applications in other heart diseases.

### Study Approval

Animal protocols were approved by the Institutional Animal Care and Use Committee of Peking University (No. DLASBD0022).

## Data availability

AAV plasmids are available at Addgene. Data, materials and methods are available upon reasonable requests.

## Acknowledgements

We thank PackGene Biotech for AAV production.

## Author Contributions

Y.G. conceived the research and wrote the paper. G.C. executed the experiments and analyzed the data. Y.Z. assisted in sample collection and analysis. Z.C. and L.Y. independently validated the finding of this study. Y.G., E.D. and F.G. supervised the study.

## Sources of Funding

This work was funded by Beijing Natural Science Foundation (7232094 to Y.G.), the National Key R&D Program of China (2022YFA1104800 to Y.G.), the National Natural Science Foundation of China (82222006 to Y.G., 82070235 to E.D. and 92168113 to E.D.) and the CAMS Innovation Fund for Medical Sciences (2021-I2M-5-003 to E.D.).

## Notes

**Conflict of interest:** The authors have declared that no conflict of interest exists.

### Competing Interest Statement

The authors have declared no competing interest.

